# Comparison of Three Individual Identification Algorithms for Sperm Whales (*Physeter macrocephalus*) after Automated Detection

**DOI:** 10.1101/2021.12.22.473895

**Authors:** Drew Blount, Jason Holmberg, Jason Parham, Shane Gero, Jonathan Gordon, J. Jacob Levenson

## Abstract

Photo-identification of individual sperm whales (*Physeter macrocephalus*) is the primary technique for mark-recapture-based population analyses for the species The visual appearance of the fluke — with its distinct nicks and notches — often serves as the primary visual differentiator, allowing humans to make recorded sightings of specific individuals. However, the advent of digital photography and the significant increase in volume of images from multiple projects in combination with pre-existing historical catalogs has made applying the method more challenging.with the required human labor for de-duplication (reduction of Type II errors) and reconciliation of sightings between large datasets too cost- and time-prohibitive. To address this, we trained and evaluated the accuracy of PIE v2 (a triplet loss network) along with two existing fluke trailing edge-matching algorithms, CurvRank v2 and Dynamic Time Warping (DTW), as a mean to speed comparison among a high volume of photographs. Analyzed data were collected from a curated catalog of well-known sperm whales sighted across years (2005-2018) off the island of Dominica. The newly-trained PIE model outperformed the older CurvRank and DTW algorithms, and PIE provided the following top-k individual ID matching accuracy on a standard min-3/max-10 sighting training data set: Rank-1: 87.0%, Rank-5: 90.5%, and Rank-12: 92.5%. An essential aspect of PIE is that it can learn new individuals without network retraining, which can be immediately applied in the presence of (and for the resolution of) duplicate individuals in overlapping catalogs. Overall, our results recommend the use of PIE v2 and CurvRank v2 for ID reconciliation in combination due to their complementary performance.

## Introduction

Historically, sperm whales (*Physeter macrocephalus*) were heavily hunted, and remain globally designated as Vulnerable on the IUCN Red List of Threatened Species (Taylor et al. 2019). Individuals can be reliably and repeatedly identified by the unique nicks and notches along their flukes (Armbom 1987; Whitehead 1990), which can then form the basis of photo-identification-based mark-recapture models to estimate population size and structure (e.g. Matthews et al. 2001; Whitehead and Gero 2015).

The Flukebook.org online platform (Blount et al. submitted; Flukebook 2021) provides a Web-based data management framework and a computer vision pipeline (Parham et al. 2018) for detection and individual identification of multiple species of cetaceans, including sperm whales. However, existing fluke matching techniques deployed on Flukebook (Jablons 2016; Weideman et al. 2020) have not yet been evaluated specifically on sperm whales, leaving questions about their accuracy and reliability for this species. Additionally, new developments in machine learning have suggested that a new class of deep learning-based algorithms may offer an improvement in automated matching capability (Capgemini 2020).

If we are going to address conservation and management for this species and conduct analyses over biologically relevant scales, beyond those of any single research group, we must be able to reconcile multiple sperm whale photo-ID catalogs with the potential for internal matches of individuals To do so, we need to automate a pipeline which is both more efficient in terms of expert time, and reduces errors of misidentification. Our team explored and analyzed three computer vision algorithms to understand their performance at matching individual sperm whales sighted across years. Here, we test three algorithms applied to this species for the first time. Two of the algorithms were Oriented Curve/Dynamic Time Warping (OC/DTW) (Jablons 2016), and CurvRank v2 (Weideman et al. 2020) both of which were off-the-shelf models previously trained on humpback whale (*Megaptera novaeangliae*) flukes. The third is called Pose Invariant Embeddings (PIE v2; Moskvyak et al. 2019), which was first tested on manta ray bellies (*Mobula* spp.) and humpback whale flukes, employs a machine learning-based triplet loss network in a fashion similar to the winning entry in a related machine learning competition exploring matchability of sperm whale flukes (Capgemini 2020). Understanding their performance can inform the usability of the Flukebook.org platform and aid researchers in not only rapidly matching individuals across photo ID catalogs (a significant potential time and cost savings on the road to larger and more comprehensive catalogs and modeling efforts) but also in understanding and potentially modeling performance-based biases in mark-recapture models.

## Materials and methods

### Dataset

We conducted our experiments and evaluations with already curated photos of well-known individuals photographed between 2005 and 2018 from The Dominica Sperm Whale Project (DSWP; Gero et al. 2014) as recorded in the Flukebook.org platform (Flukebook 2021). Our project had access to 5917 annotations (i.e., ML-detected bounding boxes around sperm whale flukes in photos) from 512 individual sperm whales.

**Figure 1.**
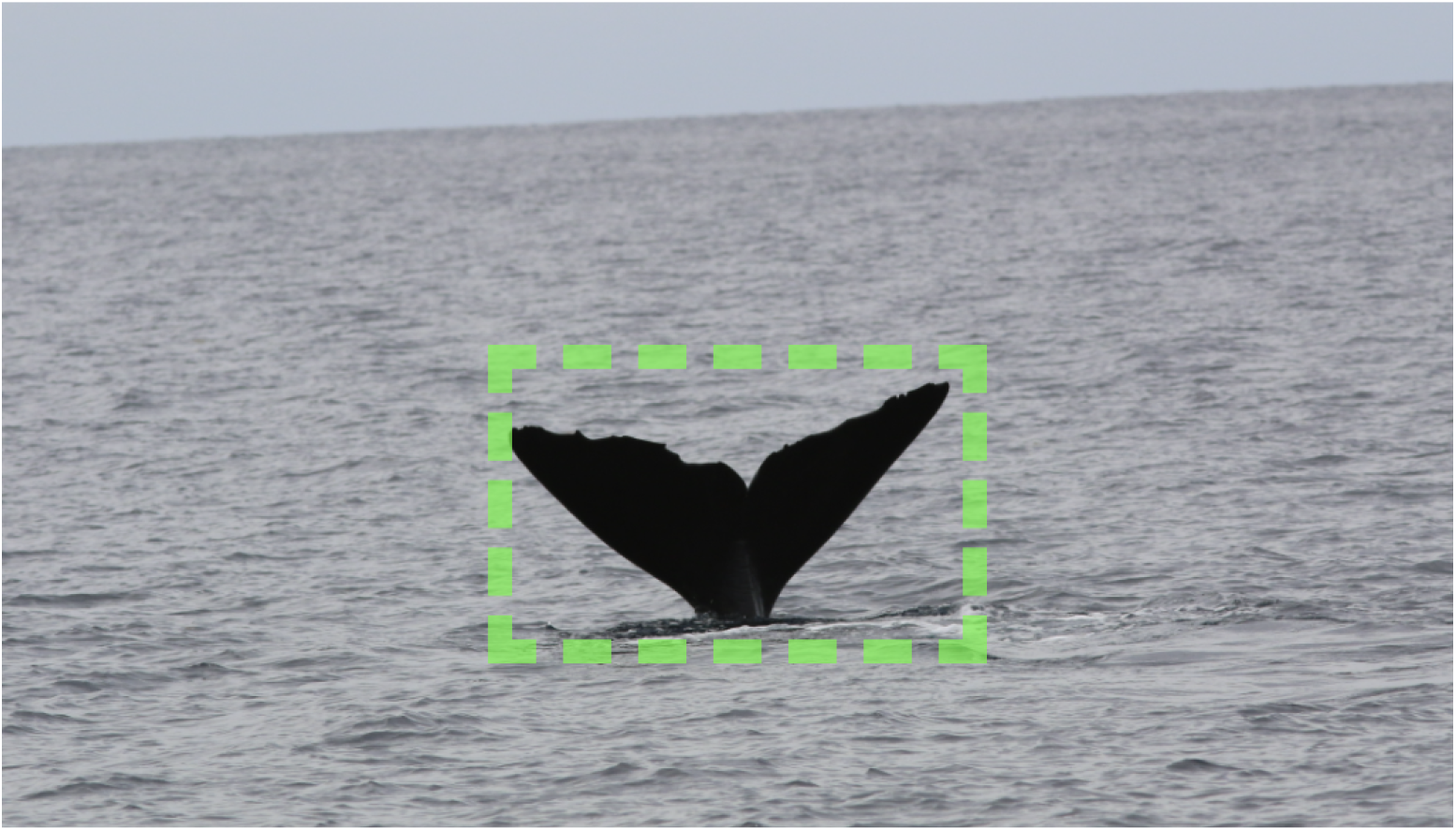
An annotated sperm whale fluke. Annotations were generated by a machine learning-based computer vision model and associated with the identification of known individuals by human confirmation. Annotations served as the fundamental data learned from by ML and compared by each algorithm. Photo courtesy of The Dominica Sperm Whale Project.

#### Performance Metric

We evaluate the performance of each algorithm individually by computing the top-k accuracy on a test set where k = 1, 5, 10 and represents the rank of the correct match (i.e. an annotation of the same individual represented by a query annotation) in a list of proposed matches. A top-1 rank therefore is the correct result returned by the algorithm as the most likely match for a candidate annotation. A top-5 rank is the correct result ranked fifth most likely as returned in the candidate list and so forth.

#### Min-3/Max-10 versus Min-2/No-max

In reviewing our performance evaluation metrics and charts, it is essential to understand which data is used and why in computing top-k performance. For example, confusion around reported matching numbers can be caused when apples-to-oranges comparisons are made between reported numbers based on differing data. Here, we use two different sets of the DSWP-source data.

#### Min-3/Max-10

For training machine learning algorithms like PIE, we often utilize a Min-3/Max-10 data subset. This report represents the subset of individuals with a minimum of three fluke annotations and a maximum of 10 allowed. A minimum of three is required for the training phase (2 photos for ML to learn from) and the test phase (at least 1 for ML to test it against). A maximum of 10 photographs is allowed for data set balance and to prevent highly sighted individuals (e.g., whale “Pinchy” from the DSWP has almost 700 encounters) from causing the ML system to optimize on highly sighted individuals yet perform poorly on infrequently sighted individuals. In our experience, a max-10 limit will suppress the Top-k performance ranking but create an ML model that performs better in real-world matching across a variety of whales. In this report, our Min-3/Max-10 data set contained 2249 annotations (i.e., ML-detected bounding boxes around sperm whale flukes in photos) from 420 individual sperm whales.

#### Min-2/No-Max

For evaluating the matching performance of one or more algorithms that have already been trained, we can relax our data constraints and require only a minimum of two annotations (i.e., we need at least two photos per individual to test matchability). We can also remove the upper limit of annotations and allow a more extensive evaluation that includes many images of frequently sighted individuals. In this case, the extended dataset used all 5917 annotations from 1,188 individual sperm whales provided by DSWP.

### Fluke Detection

Before the algorithms can act, sperm whale flukes need to be detected in the images. Sperm whale fluke annotations were created by a machine learning detector in the WBIA pipeline (Parham et al. 2018). The task of the detector — a customized PyTorch implementation of YOLO v2 (Redmon et al. 2016) used to localize animals in images — is focused mainly on localizing accurate bounding boxes over the ground-truth detections (made *a priori* by humans for a test set) while minimizing false positives and false negatives. We trained a model to predict the whale_sperm+fluke class using a training dataset of 1,592 images and 1,427 annotated bounding boxes (184 empty images, 19 images with two boxes).

Figure 2 shows the Precision-Recall performance curves of the trained sperm whale detection model as computed on a held-out 20% test set (286 annotations). The various colors show different thresholds of non-maximum suppression (NMS) applied to the network’s final bounding box predictions. A common way to summarize the localization accuracy is with Average Precision (AP) as determined by the area-under-the-curve. For example, the best performing configuration with an NMS of30% achieves an AP of 91.65%. The corresponding colored points on each curve signify the closest point along the line to the top-right corner of the precision-recall coordinate system, signifying a perfect detector. Furthermore, the yellow diamond specifies the highest precision for all configurations, given a desired fixed recall of 80%. The confusion matrices below in Figure 3 give the accuracy for the best-colored point (left) and the yellow diamond (right). It is worth mentioning that the 80% recall is arbitrary and can be adjusted based on the performance targets of the final project.

**Figure 2.**
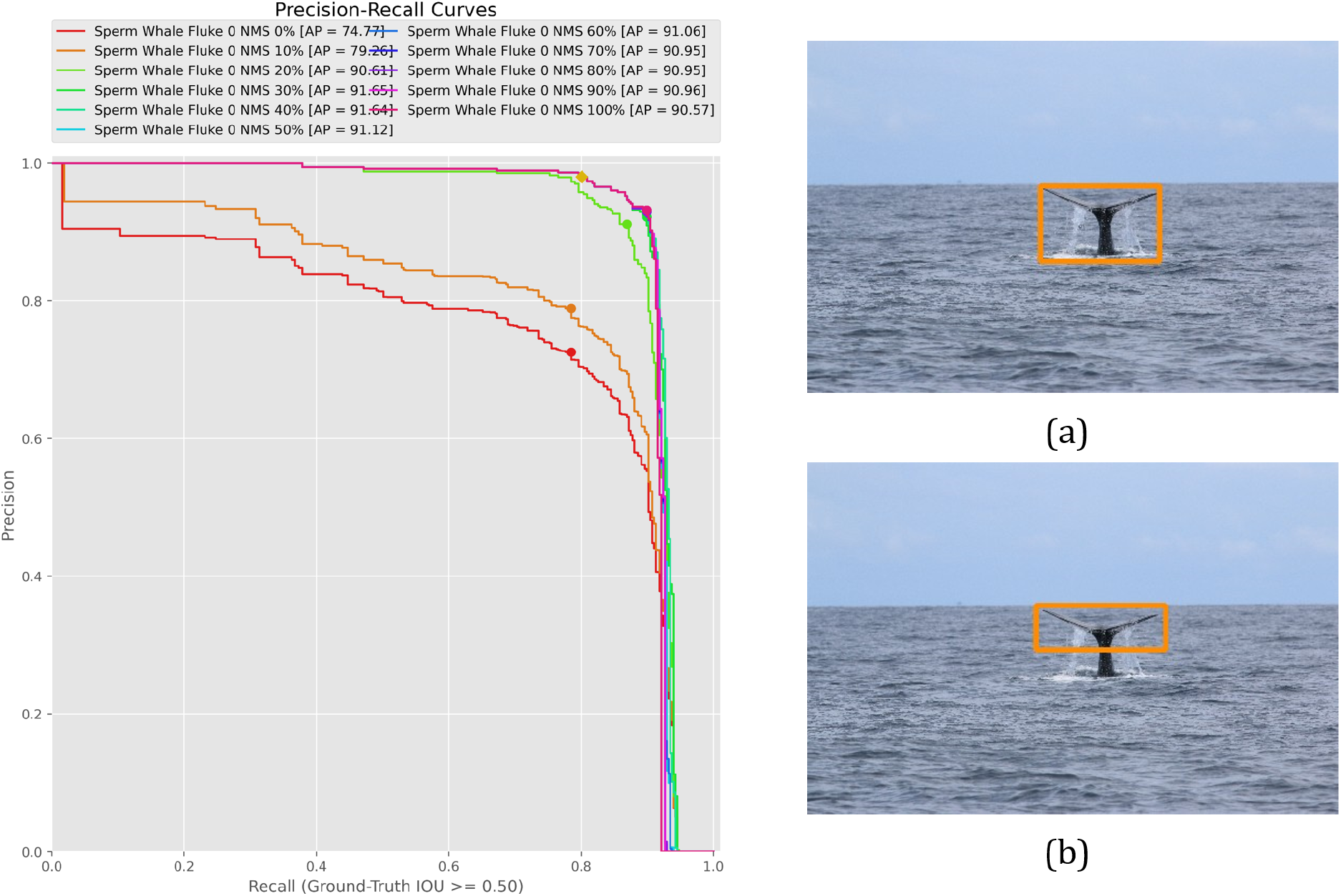
The detector Precision-Recall curves for sperm whale flukes. We also show an example held-out test image with an actual detection (b) against the ground-truth bounding box (a). The bounding boxes are produced with NMS=40% and operating point of 65%.

**Figure 3.**
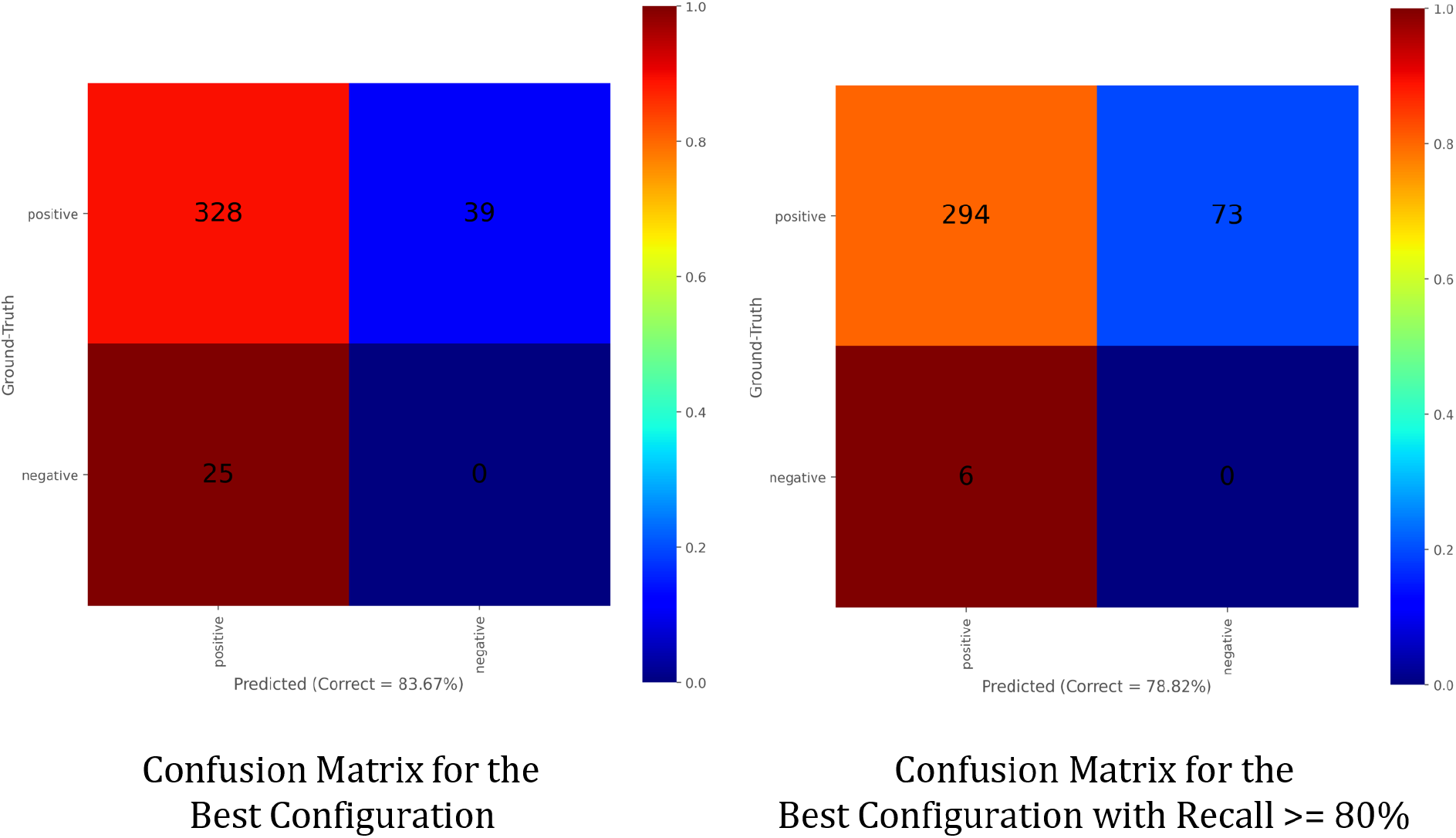
Confusion matrices for the best-colored point (left) and the yellow diamond (right) from the Precision-Recall performance curves in Figure 2. False negatives occur when not detecting a fluke when one is present in the image and false positives are spurious detections when no fluke is present in the image, such that a bounding box is generated where there is no fluke within it.

Delving deeper on the Precision-Recall curves, the maximum recall values (x-axis intercept) represent the absolute maximum percentage of annotations that the detector configuration can “recover” or “recall” from the ground-truth detections. Therefore, a recall of 90% indicates that a given detection configuration found 90% of the ground-truth annotations. The recall is a fundamental measurement for false negatives and implies a miss rate of 10% per sighting. The precision value indicates the percentage of correct detections (and thereby also providing a measure for the number of false positives) and how many additional incorrect detections are given. A true-positive in our detection scenario is defined by the amount of intersection-over-union (IoU) percentage between a prediction and a matched ground-truth bounding box. For all plots in this section, we fix the acceptable IoU threshold to be 50% or greater. Non-maximum suppression is a common technique for filtering duplicate detections by eliminating highly overlapping and lower-scoring predictions. A high NMS value will remove many bounding boxes from the output based on their percentage of overlap area (leading to an increase in precision but a decrease in recall). True negatives are undefined, which is why a receiver operating characteristic (ROC) curve is not provided in this report.

Looking closely at Figures 2 and 3, we can see that the best performing and our chosen configuration (highest AP at over 90%) has an NMS threshold of 30% and a score threshold of 67%. With this configuration, the overall detector makes 64 errors out of 367 overall annotations, with 39 of those incorrect detections being false negatives (not detecting a fluke when one is present). Figure 3 shows that for 39 false negatives, there are 25 false positives (spurious detections of flukes that were not in the image, a bounding box is generated where there is no fluke). If we relax the miss-rate requirement to 20%, we make fewer false detections (a total of 6 down from 25), but we end up missing 73 animals (false negatives), and the overall accuracy drops by nearly 5% (see Figure 3, right plot).

### Network architecture, data preprocessing, training, and evaluation

With a machine learning detector trained and configured to extract sperm whale fluke annotations, we then used those annotations and related metadata (in particular the known identifications of the flukes) in the WBIA pipeline (Parham et al. 2018) to first custom train the Pose Invariant Embeddings (PIE) algorithm (Moskvyak et al. 2019) and then evaluate a) the custom-trained PIE model, b) the pre-trained CurvRank v2 algorithm (Weideman et al. 2020; trained on humpback flukes), and c) the pre-trained OC/DTW (Jablons et al. 2016) algorithms. The CurvRank v2 and OC/DTW algorithms had already been deployed for sperm whale fluke matching in Flukebook.org and shown anecdotally to be able to match individuals based on their independent extraction of the trailing edge of the fluke. While OC/DTW cannot be custom-trained in its current implementation, our choice to cross-apply the existing CurvRank v2 model trained originally on annotated humpback whale flukes was deliberate, allowing us to contrast a known production model already in use on Flukebook.org with the custom training of the triplet-loss-based PIE model, which does not extract a fluke but rather learns from all the pixels of an annotation. PIE represents a novel application of a new machine learning architecture as suggested by the unpublished results of a recent machine learning competition (Capgemini 2020).

## Experiments (PIE training)

We used a Min-3/Max-10 data constraint for PIE ML training and divided the training and test data according to Table 1.

**Table 1.** Data division for PIE machine learning training.

| Set | Individuals | Annotations |
| --- | --- | --- |
| train | 310 | 1587 |
| test | 110 | 652 |
| total | 420 | 2249 |

## Results

On the required Min-3/Max-10 data basis, our top-performing PIE model returned top-k ranks of:

- Top-1: 87.0%
- Top-5: 90.5%
- Top-12: 92.5%

Because Flukebook contains a multi-species, multi-feature, and multi-algorithm technical foundation (Flukebook 2021; Blount et al. Submitted), more than one algorithm can be run in parallel when identifying the individual animal in a photo. Therefore to evaluate PIE v2, Dynamic Time Warping, and CurvRank v2, we used our Min-2/No-Max data set and evaluated and plotted all algorithms and their combinations to visualize their relative strengths and to suggest an optimal algorithm combination for deployment. Figure 4 shows their relative performances.

**Figure 4:**
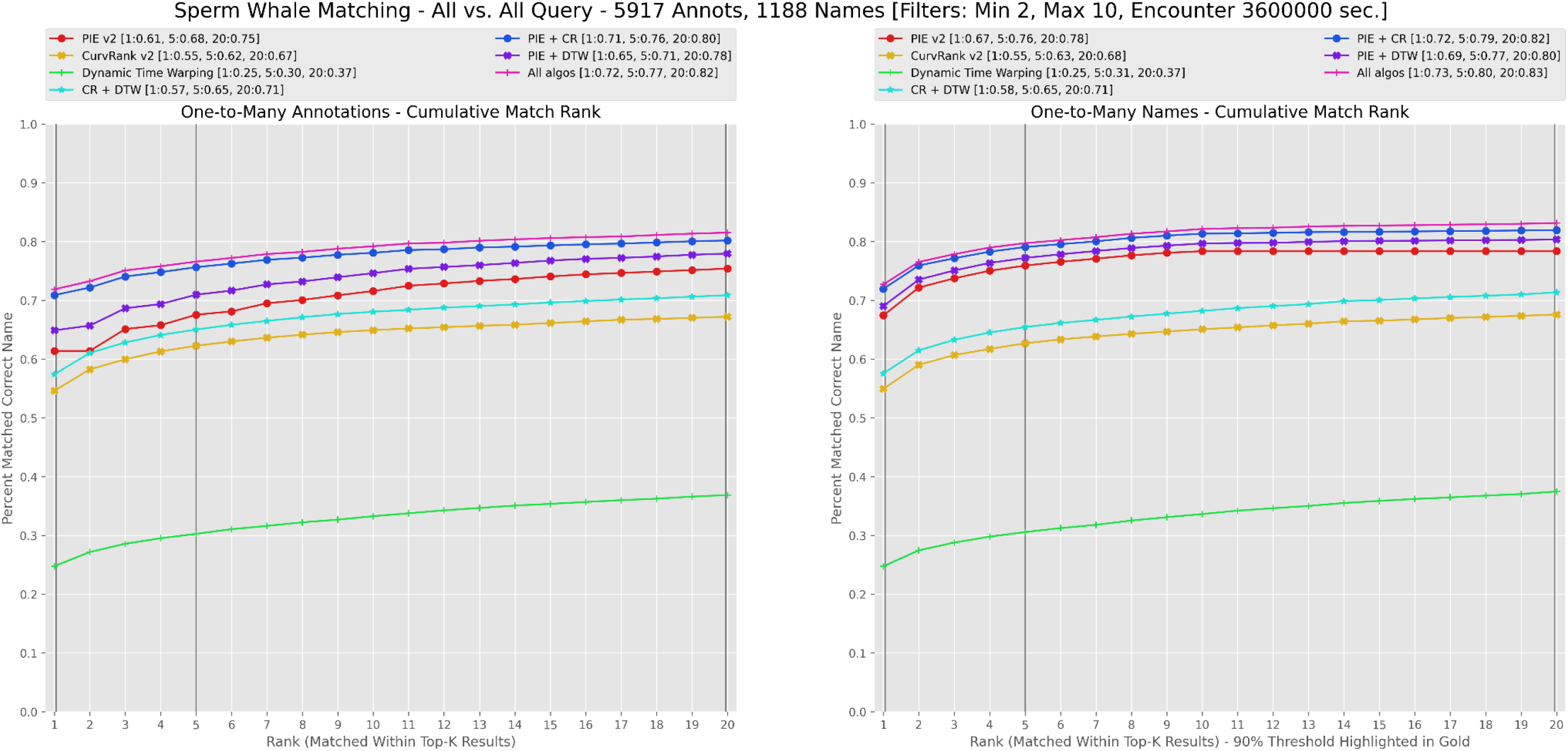
Matching accuracy of PIE v2, CurvRank v2, OC/DTW, and any combination thereof with Min-2/No-Max data. On the left is annotation-matching, which is the accuracy of finding a single candidate annotation with the correct individual name assuming only that match for the name exists in the query set. The right-side displays name-matching (finding any annotation of the individual at the listed rank), which is the accuracy at assigning the correct name to a query annotation wherein 1 or more matches may be available for a name. Combined accuracies represent how often any of the listed algorithms return the same ID result at rank k.

Among the evaluated techniques, the PIE v2+CurvRank v2 algorithms provided the most overall individual matching power with an additive performance of top-1 rank of 70% and top-12 of 82%. This dataset represents the largest and most challenging collection available for analysis, meaning these numbers are highly robust, and higher accuracy can be expected when filtering to smaller datasets. The PIE model outperformed the older CurvRank and DTW algorithms, and PIE provided the following top-k individual ID matching accuracy on a standard min-3/max-10 sighting training data set: Rank-1: 87.0%, Rank-5: 90.5%, and Rank-12: 92.5% (Figure 4).The large dataset size with a minimum of two sightings/individuals explains the discrepancy between the former accuracy numbers and these min-3 accuracy.

## Discussion

Our results provide a first look at not only the matching accuracy of three independent, production-ready fluke-matching algorithms for sperm whales but also demonstrate that their use in combination can further improve researchers’ ability to rapidly identify matching flukes across large catalogs. The highest performing algorithm, PIE, has generally been considered a pattern-matching algorithm (Moskvyak et al. 2019), but being a deep learning technique, it has remained somewhat of a black-box in terms of which features it uses for matching decisions. Our efforts here, and those previously of Capgemini (2020), represent very new applications of emerging ML on a non-patterned species, and the results are very encouraging. There is little-to-no identifiable pattern on the all-black fluke of a sperm whale, so the fact that PIE has outperformed more traditional trailing-edge matching algorithms is significant. This high performance suggests that the pattern-matching abilities of the PIE network are flexible enough to effectively distinguish the trailing edge of the fluke from background water or sky and use that area for matching. The pattern of PIE outperforming CurvRank and OC/DTW is likely to be seen for other contour-based species. In the future, we intend to train PIE on other trailing-edge-focused problems such as the dorsal fins of toothed and baleen whales, dolphins, sharks, and possibly elephant ears. This work has shown that PIE is a more general-purpose matching algorithm than was initially assumed, and at least, in this case, it out-competes specialized trailing-edge approaches in their problem domain.

Based on the comparative results presented above, we recommend a default deployment configuration of PIE v2 and CurvRank v2 for matching sperm whales in Flukebook.org, and these are now immediately available for use on the platform. PIE and CurvRank offer significant complementary performance (i.e., each can significantly catch matches missed by the other). In contrast, DTW provides no significant matching not otherwise represented in the results of PIE or CurvRank.

## Data Availability

Research-related requests for annotations and data used for ML training in this paper can be requested in COCO format (Lin et al. 2020) via the corresponding author and must be expressly and independently permitted by author Shane Gero or through an established collaboration on Flukebook.org. Data can also be reviewed and shared via a Collaboration request to user Shane Gero inside the Flukebook.org system.

## Code Availability

All software used in this analysis is available in the Wild Me open source repository at: https://github.com/wildmeorg

The base application for algorithm analysis as defined in Parham et al. 2018 is: https://github.com/WildMeOrg/wildbook-ia

Specific algorithm plugins for the three algorithms evaluated here can be found at:

- https://github.com/WildMeOrg/wbia-tpl-curvrank-v2
- https://github.com/WildMeOrg/wbia-plugin-pie-v2
- https://github.com/WildMeOrg/wbia-plugin-flukematch

## Acknowledgements

The authors would like to thank collaborator Dr. Jonathan Gordon for suggesting they review the competition-winning work of Capgemini (Capgemini 2020) in identifying individual sperm whales using an Azores-based data set from Dr. Lisa Steiner. Funding was provided in part by the U.S. Department of the Interior, Bureau of Ocean Energy Management, through Cooperative Agreement M20AC10010, sub-awardee Wild Me (wildme.org; sub-award DI132A-A) under principal investigator, C.S. Baker, Marine Mammal Institute, Oregon State University. Field research for The Dominica Sperm Whale Project over those years was funded through a FNU fellowship for the Danish Council for Independent Research supplemented by a Sapere Aude Research Talent Award, a Carlsberg Foundation expedition grant, a grant from Focused on Nature, and a CRE Grant from the National Geographic Society to SG; a FNU Large Frame Grant and Villum Foundation Grant to Peter T. Madsen at Aarhus University; NSERC Discovery and Equipment grants to Hal Whitehead; and supplementary grants from the Arizona Center for Nature Conservation, Quarters For Conservation, the Dansk Akustisks Selskab, Oticon Foundation, and the Dansk Tennis Fond.

## Author contributions

AB co-wrote and performed the PIE algorithm training and multi-algorithm analyses. JH co-wrote, fundraised and coordinated data annotation for the project. SG collected and curated the data set used for machine learning training and performance analysis. JG provided insight into photo ID challenges and related efforts. JL provided administrative support. All authors provided comments and final approval of the uploaded manuscript.

